# The Effects of Sex and Diet on Physiology and Liver Gene Expression in Diversity Outbred Mice

**DOI:** 10.1101/098657

**Authors:** Daniel M. Gatti, Petr Simecek, Lisa Somes, Clifton T. Jeffrey, Matthew J. Vincent, Kwangbom Choi, Xingyao Chen, Gary A. Churchill, Karen L. Svenson

**Affiliations:** The Jackson Laboratory, Bar Harbor, ME 04609, USA

**Keywords:** Diversity Outbred, Nutrigenomics, QTL, Complex Traits

## Abstract

Inter-individual variation in metabolic health and adiposity is driven by many factors. Diet composition and genetic background and the interactions between these two factors affect adiposity and related traits such as circulating cholesterol levels. In this study, we fed 850 Diversity Outbred mice, half females and half males, with either a standard chow diet or a high fat, high sucrose diet beginning at weaning and aged them to 26 weeks. We measured clinical chemistry and body composition at early and late time points during the study, and liver transcription at euthanasia. Males weighed more than females and mice on a high fat diet generally weighed more than those on chow. Many traits showed sex- or diet-specific changes as well as more complex sex by diet interactions. We mapped both the physiological and molecular traits and found that the genetic architecture of the physiological traits is complex, with many single locus associations potentially being driven by more than one polymorphism. For liver transcription, we find that local polymorphisms affect constitutive and sex-specific transcription, but that the response to diet is not affected by local polymorphisms. We identified two loci for circulating cholesterol levels. We performed mediation analysis by mapping the physiological traits, given liver transcript abundance and propose several genes that may be modifiers of the physiological traits. By including both physiological and molecular traits in our analyses, we have created deeper phenotypic profiles to identify additional significant contributors to complex metabolic outcomes such as polygenic obesity. We make the phenotype, liver transcript and genotype data publicly available as a resource for the research community.

## INTRODUCTION

Many factors affect the physiology and transcriptional landscape of individuals. Intrinsic factors, such as sex and genetic background, play a role in shaping individuals. External factors, such as diet and other environmental stimuli, also play a role in determining the health and well-being of each person. However, wide variation in individual response to diet is an enormous challenge to developing prevention and treatment strategies aimed at reducing the incidence of obesity and metabolic disorders. The response to diet is affected by both sex (Griffin *et al.* 2016) and individual genetic background (Suhre and Gieger 2012). Disparity in obesity prevalence among sexes has been ascribed to both biological and sociocultural factors (Kanter and Caballero 2012). Sex differences in fat gain and storage can lead to considerable differences in health outcome among women and men (Power and Schulkin 2008). Additionally, sex-dependent single nucleotide variants have been reported that underlie differential contributions to the development of obesity (Kvaloy *et al.* 2013; Saldana-Alvarez *et al.* 2016).

In human populations, it is difficult to dissect the genetic basis for differential responses to diet between the sexes due to differences in lifestyle and uncontrolled covariates. While epidemiological models can estimate correlations between different traits, the controlled conditions in mouse models allow us to apply randomization and factorial designs to detect causal associations between obesity and physiological traits. In mouse models, we can also control the genetic background of the mice, thereby reducing another uncontrolled variable that influences the response to diets between the sexes. Most mouse models of obesity use a single inbred strain genetically engineered or experimentally manipulated to become obese (reviewed in (Hariri and Thibault 2010)). However, genetic background is known to influence the response of individuals to dietary fat (Wang *et al.* 2002; Stoehr *et al.* 2004; Su *et al.* 2008; Lin *et al.* 2013). In order to generalize our results from mice to humans, it is critical to include structured genetic diversity in mouse models of dietary response.

Multi-parent advanced intercross (MAGIC) populations are powerful models for mapping genetic modifiers of complex traits due to their high minor allele frequency and fine mapping resolution (Churchill *et al.* 2004; Rakshit *et al.* 2012; Rat Genome *et al.* 2013; Gatti *et al.* 2014). In MAGIC populations derived from known founders, the haplotype structure of each sample genome can be reconstructed in terms of the founder genomes (Mott *et al.* 2000; Gatti *et al.* 2014). When the founders have been fully sequenced, the founder sequences can be imputed onto the MAGIC genomes to allow for whole genome association mapping (Yalcin *et al.* 2005), which improves the ability to identify candidate genes that influence traits. Transcript profiling in a relevant tissue adds another important dimension to genetic mapping studies and can be used to perform mediation analysis on each significant genomic locus (Chick *et al.* 2016). There are several mouse MAGIC populations available, including the Northport Heterogeneous Stock (Valdar *et al.* 2006), the Collaborative Cross (Threadgill and Churchill 2012), the Heterogeneous Stock/Collaborative Cross (Iancu *et al.* 2010) and the Diversity Outbred (Svenson *et al.* 2012).

In this study, we fed Diversity Outbred mice of both sexes either standard chow or a high-fat/high-sucrose diet from weaning until approximately 6 months of age. We measured a variety of physiological traits throughout the study and performed liver transcriptional profiling at the end of the study. Here, we report on the differential effects of diet on each sex in terms of physiological and transcriptional trait, provide interactive viewers for the results and release the entire data set to the public through supplemental materials and at http://do.jax.org.

## METHODS

### Mice and husbandry

Diversity Outbred mice were obtained from The Jackson Laboratory (Bar Harbor, ME). This study used five independent cohorts of 100-200 non-sibling DO mice from generations 4 to 11 (G4-G11) for a total of 850 animals, which builds on an initial study that has previously been reported (Svenson *et al.* 2012). In each cohort, half the animals were from first litters in the respective generation and half were from second litters. An equal number of females and males were included in each set of animals received. Mice were housed at a density of five same-sex mice per pen in pressurized, individually ventilated cages (Thoren #11 Duplex II; Thoren Caging Systems, Hazelton, PA) with pine bedding (Crobb Box, Ellsworth, ME) and free access to food (diets described below) and acidified water. Light cycle was 12h:12h light:dark, beginning at 0600. All animal procedures were approved by the Animal Care and Use Committee at The Jackson Laboratory (Animal Use Summary # 06006).

### Phenotyping

Upon receipt, when mice were 3 weeks of age, equal numbers of each sex were randomly assigned to chow (LabDiet 5K52, LabDiet, Scott Distributing, Hudson, NH) or high fat, high sucrose feeding (Envigo Teklad TD.08811, Envigo, Madison, WI) for the duration of the study protocol (26 weeks). Caloric content of the high fat diet (HFD) was 45% fat, 40% carbohydrates and 15% protein. Tail biopsies were taken at wean for DNA preparation. Weight was monitored weekly throughout the study. At age 8 weeks mice began a pipeline of noninvasive phenotyping assays to profile metabolic health (Table 1). Some modifications to the pipeline were made as the number of cohorts progressed, such that all parameters were not measured in all mice. Table 1 lists each phenotypic measurement and the number of mice tested. Data was obtained from 846 mice and 154 traits were used for analysis. Clinical chemistries, urinalysis and body composition assessments were performed at two time points in the study to evaluate stability of traits under prolonged HFD. Hence, calculated traits comparing first and second measures were generated as derived traits. Details about blood collection and analysis and body composition by dual-energy x-ray absorptiometry (DEXA) have been described previously for this pipeline (Svenson 2012 Genetics). Additional tests include body composition by qNMR (EchoMRI), electrocardiogram, intraperitoneal glucose tolerance test (ipGTT), and evaluation of chemokines by electrochemiluminescence. To quantitate lean and fat tissue and free and total water, EchoMRI (EchoMRI, Houston, TX) without anesthesia was used, providing three time points for evaluation of tissue composition during the study and minimizing the need for anesthesia during the pipeline. Electrocardiography was performed using the ECGenie™ (Mouse Specifics, Quincy, MA) system, whereby unanesthetized mice are placed on a platform raised 18” above the laboratory bench containing a lead plate. When animals contact the plate with any three paws the trace begins. Fast Fourier analysis (AnonyMouse™ software v2.2; Mouse Specifics) defines interval durations from which heart rate, variability and other features of cardiac conduction can be assessed. To evaluate glucose clearance, mice were fasted overnight (15 hours) and in the morning mice were weighed and a small blood sample from a tail tip incision was used in the Abbott glucometer system to measure fasted glucose (GTT time 0; t0). A glucose solution was then injected intraperitoneally at 2 mg glucose/gram body weight and tail tip blood samples were obtained at 15, 30, 60, 120 and 180 minutes after injection. Plasma leptin, insulin, ghrelin and adiponectin were measured from nonfasted mice using the Meso Scale Discovery™ electrochemiluminescent assay detection system according to the manufacturer’s protocols. We transformed all traits to ranked Z-scores before performing statistical analyses.

### Genotyping and Diplotype Reconstruction

DNA was prepared from tail biopsies and genotyped using two versions of the Mouse Universal Genotyping Array (MUGA) (Morgan *et al.* 2016). We genotyped 531 samples on the MUGA and 293 samples on the Megamuga (GeneSeek, Lincoln, NE). We used the intensities from each array to infer the haplotype blocks in each DO genome using a hidden Markov model (HMM) (Gatti *et al.* 2014).

### Genotyping by RNA-sequence

Genotyping by RNA-sequence (GBRS) is a set of software tools that reconstruct individual genomes of each sample in multi-parent population (MPP) models by decoding known polymorphisms of founder strains from RNA-Seq data without resorting to genotyping arrays. The new method is efficient since it avoids maintaining hundreds of individualized genome indexes by aligning RNA-Seq reads to a common pooled transcriptome of all founder strains a single time. Since our reusable model parameters can be easily estimated from separate RNA-Seq data of inbred founder strains or from simulations, we can quickly process each MPP sample independently. The software package implements our alignment strategy and statistical models and is freely available at https://github.com/churchill-lab/gbrs. We used the GBRS haplotype reconstructions to fill in samples that failed to genotype due to low call rates on the MUGA or Megamuga.

### Merging Haplotype Reconstructions from Different Methods

The MUGA and Megamuga have different numbers of markers (MUGA: 7,854, Megamuga: 77,642) and the HMM produced diplotype probabilities only at each marker. In contrast, the GBRS method produced diplotype probabilities at each gene that was expressed in the liver. In order to merge diplotype probabilities from the data, we interpolated both markers grids to an evenly spaced 64,000 marker grid (0.0238 cM between markers). After merging the diplotype reconstructions, we had a total of 835 samples.

### Principal Component Analysis of Physiological Traits and Liver Transcription

We retained 129 out of 160 physiological traits with < 50% missing data across samples. The 24 traits with > 50% missing data were ACR1, ACR2, Adiponectin, BW.3, BW.27, BW.28, BW.29, BW.30, fat.g.mri, free.h20, FRUC1, Ghrelin, GTT.AUC, GTT.t0, GTT.t15, GTT.t30, GTT.t60, GTT.120, GTT.180, Lipase1, non.fast.Calcium, TBIL1, TBIL2 and tot.h20 (see File S1 for abbreviations). We used the Probabilistic PCA method of the pcaMethods software (Stacklies *et al.* 2007) to impute missing data in the remaining traits and calculated the first 10 principal components.

### Physiological Traits Correlations

We regressed out the outbreeding generation, sex and diet effects from each of the physiological traits and calculated the pairwise Pearson correlation between all physiological traits.

### Alignment, Quantification and Normalization of Liver Transcription Data

We aligned reads from the DO liver data to pooled transcriptomes derived from the eight DO founder strains by incorporating strain-specific SNPs, insertions and deletions into the reference genome sequence. We quantified expected read counts using an expectation maximization algorithm (EMASE, https://github.com/churchill-lab/emase) (Chick *et al.* 2016). We retained 12,067 genes with mean transcripts per million across all samples greater than one. We normalized effective counts to the upper quartile value and transformed them to rank normal scores.

### Differential Expression and Gene Set Enrichment Analysis of Liver Transcript Data

We performed analysis of variance (ANOVA) on the normalized liver transcription data to identify genes that were differentially expressed between sexes, diets or that had a sex by diet interaction. We regressed the expression of each gene on generation and litter, sex, diet and the sex by diet interaction. We adjusted the p-values using the Benjamini & Hochberg false discovery rate (FDR)(Benjamini and Hochberg 1995).

We searched for Gene Ontology (GO) categories (Ashburner *et al.* 2000) that were differentially expressed between sexes, diets or that had a sex by diet interaction using the SAFE software package (Barry *et al.* 2005). We used the “t.Student” local statistic to test for differential expression for each gene and the “Wilcoxon” global statistic to test for differential enrichment between categories. For sex effects, we regressed diet from each gene and then tested for the effect of sex. For diet effects, we regressed sex from each gene and then tested for the effect of diet. For the sex by diet interaction, we compared the reduced model with sex and diet to the full model containing sex, diet and the sex by diet interaction. We determined the empirical p-value for each category using 10,000 permutations and retained GO categories with p-values ≤ 0.05.

### Linkage Mapping

At each marker, we regressed each phenotype on generation, sex, diet and the diplotype probabilities for each mouse and included an adjustment for correlation between residuals due to kinship. We used the same model for liver expression QTL mapping. We performed 5,000 permutations of a rankZ transformed phenotype and selected significance thresholds from the empirical distribution of maximum LOD scores. We estimated the founder allele effects using a Best Linear Unbiased Predictor (BLUP) in which we fit a mixed-effects model at each marker that shrinks the founder effects in proportion to the magnitude of the standard errors. We used the *qtl2* R package available at: https://github.com/rqtl. Full details of the linkage mapping model are in (Gatti *et al.* 2014).

### Association Mapping

We imputed the DO founder SNPs from the Sanger Mouse Genomes Project (REL-1505) onto each founder haplotype block in the DO genomes. We then regressed each phenotype on generation, sex, diet and the SNP probabilities for each mouse and included an adjustment for correlation between residuals due to kinship. Full details of the association mapping model are in (Gatti et al. 2014).

### Mediation Analysis

For each physiological QTL peak with a genome-wide adjusted p-value above 0.05, we performed mediation analysis to identify candidate liver genes that might be responsible for the peak (Chick *et al.* 2016). We fit a null model by regressing the phenotype on generation, sex, diet and the diplotype probabilities at the markers with the maximum LOD score. We added the expression of each of the 12,067 genes to the model and recorded the drop in the LOD score compared to the null model. We estimated the standard deviation of the LOD drops and report genes that decreased the LOD score by more than 6 standard deviations. We refer to these standardized LOD score drops as “Z-scores”. We used the *intermediate* R package available at https://github.com/simecek/intermediate.

### Data and Reagent Availability

J:DO mice are available for purchase from The Jackson Laboratory (Strain # 009376) at https://www.jax.org/strain/009376. The liver gene expression data is archived at the Short Read Archive under project number PRJNA35625. The physiological phenotypes are described in File S1, the raw phenotypes are in File S2 and the normalized phenotypes are in File S3. The genotype data for all mice and the R data objects used in all analyses are available at http://do.jax.org. We used the Sanger REL-1505 SNPs and structural variants (Keane *et al.* 2011) and the Ensembl build 82 transcripts (Yates *et al.* 2016).

## RESULTS

### Impact of Sex and Diet on Physiological Traits and Liver Transcription

We maintained mice of both sexes on either a chow diet (n = 449) or a high fat diet (n = 397) from wean to at least 26 weeks. We measured a range of physiological traits throughout the study and measured several traits at two time points (File S1). We calculated the principal components of the physiological traits and found that the mice grouped by sex and diet (Figure 1A). Principal component (PC) 1 accounted for 29.2% of the variance and is correlated with sex. PC2 accounts for 7.6% of the variance and is correlated with diet. We also quantified liver transcription at 26 weeks in a subset of 478 mice. When we calculated the PCs for a subset of 12,067 genes, we found that the samples also clustered by sex and diet (Figure 1B). PC1 and 2 accounted for 12.1% and 10.9% of the variance, respectively. Mice on the chow diet formed tighter clusters than mice on the high fat diet, reflecting larger variance in liver gene expression in mice on the high fat diet.

**Figure1.**
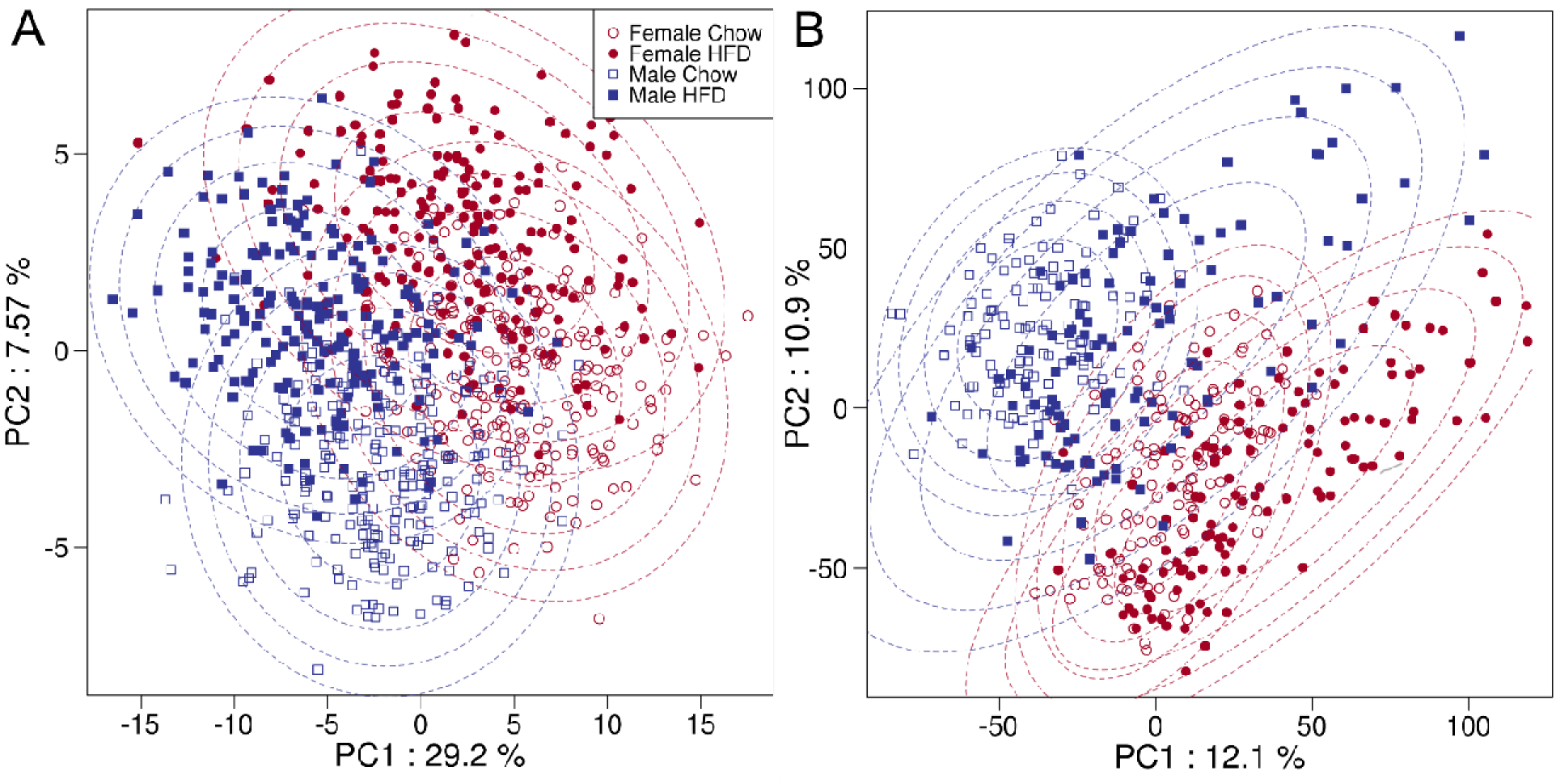
Principal component analysis (PCA) of physiological and liver transcription traits. (A) PCA plot of the first and second principal components of the physiological traits. Each point represents one mouse. Females are red; males are blue; mice on the chow diet are shown with open symbols; mice on the HFD with closed symbols. Dashed lines are ellipses from bivariate Gaussian distributions fit over each of the four sex and diet groups. (B) PCA plot of the first and second principal components of the liver transcription traits.

### Correlation between Physiological Traits

We calculated the pairwise Pearson correlation of all traits after regressing out sex and diet and identified several clusters of correlated traits (File S4). Most traits measured at two time points clustered near each other, indicating that genetic background affects many traits throughout the mouse’s lifespan. Body weight (BW) at all time points was positively correlated with other BW as well as adiponectin, insulin, bone mineral density (BMD) and lean and fat tissue mass.

Cholesterol (CHOL) at 19 weeks was highly correlated with high-density lipoprotein (HDLD, ρ = 0.95, p < 10^−16^),as expected for mice, triglycerides (TG, ρ = 0.40, p < 10^−16^), glucose (ρ = 0.28, p = 1.7 × 10^−16^), non-esterified fatty acids (NEFA, ρ = 0.44, p < 10^−16^), body weight (ρ = 0.20, p < 6.6 × 10^−9^) and circulating calcium (Ca, r = 0.50, p < 10^−16^). These correlations are in agreement with recent evidence that circulating calcium levels are associated with worsening lipid profiles in humans (Gallo *et al.* 2016) and that coronary artery calcification is an independent risk factor for atherosclerosis and cardiovascular disease (Budoff *et al.* 2007). The area under the curve of the glucose tolerance test at 24 weeks (GTT) was positively correlated with BW at 24 weeks (ρ = 0.29, p = 4.70 × 10^−5^) and other time points, GLUC at 19 weeks (ρ = 0.30, p = 1.87 × 10^6^), leptin at 8 weeks (ρ = 0.21, p = 2.62 × 10^−3^), indicating a connection between appetite and circulating glucose levels. Leptin (ρ = 0.80, p < 10^−16^), insulin (ρ = 0.42, p < 10^−16^), adiponectin (ρ = 0.40, p < 10^−16^) and % fat at both time points were positively correlated, indicating a connection between appetite and adiposity. Glutamate dehydrogenase (GLDH) at 19 weeks was positively correlated with and BW traits at ages over 15 weeks (ρ = 0.212, p = 2.45 × 10^−9^). While this observation may suggest that liver injury is associated with increased weight, it is not correlated with increased fat tissue mass as might be expected. It is likely that this correlation is an effect of aging and may be driven by those animals that were fed HFD. The correlations are available as an interactive online tool at: http://churchill-lab.jax.org/www/Svenson850/corr.html.

### Impact of Sex and Diet on Physiological Traits

We tested each trait for the effect of sex, diet and a sex by diet interaction in order to identify the effects of each on the physiological traits (File S5). This analysis stratified our results into four effect classes, each demonstrated by examples in Figure 2. There were 12 traits for which no difference was found between sexes or diet groups, including eosinophil counts (EOS) and spleen weight (Figure 2A). Sex had an effect on 130 traits at a false discovery rate (FDR) ≤ 0.05 (Figure 2B). Males had higher mean values for 101 of these traits, including body weight, monocyte counts (MONO), neutrophil counts (NEUT), glucose tolerance test (GTT.AUC), heart rate (HR), mean corpuscular volume (MCV), mean platelet volume (MPV), phosphorous, platelet counts (PLT), red blood cell distribution width (RDW) and cholesterol (CHOL). Females had higher mean values for 29 traits, including mean corpuscular hemoglobin concentration (CHCM), hemoglobin (HGB), ghrelin, %Fat and red blood cell counts (RBC).

**Figure 2.**
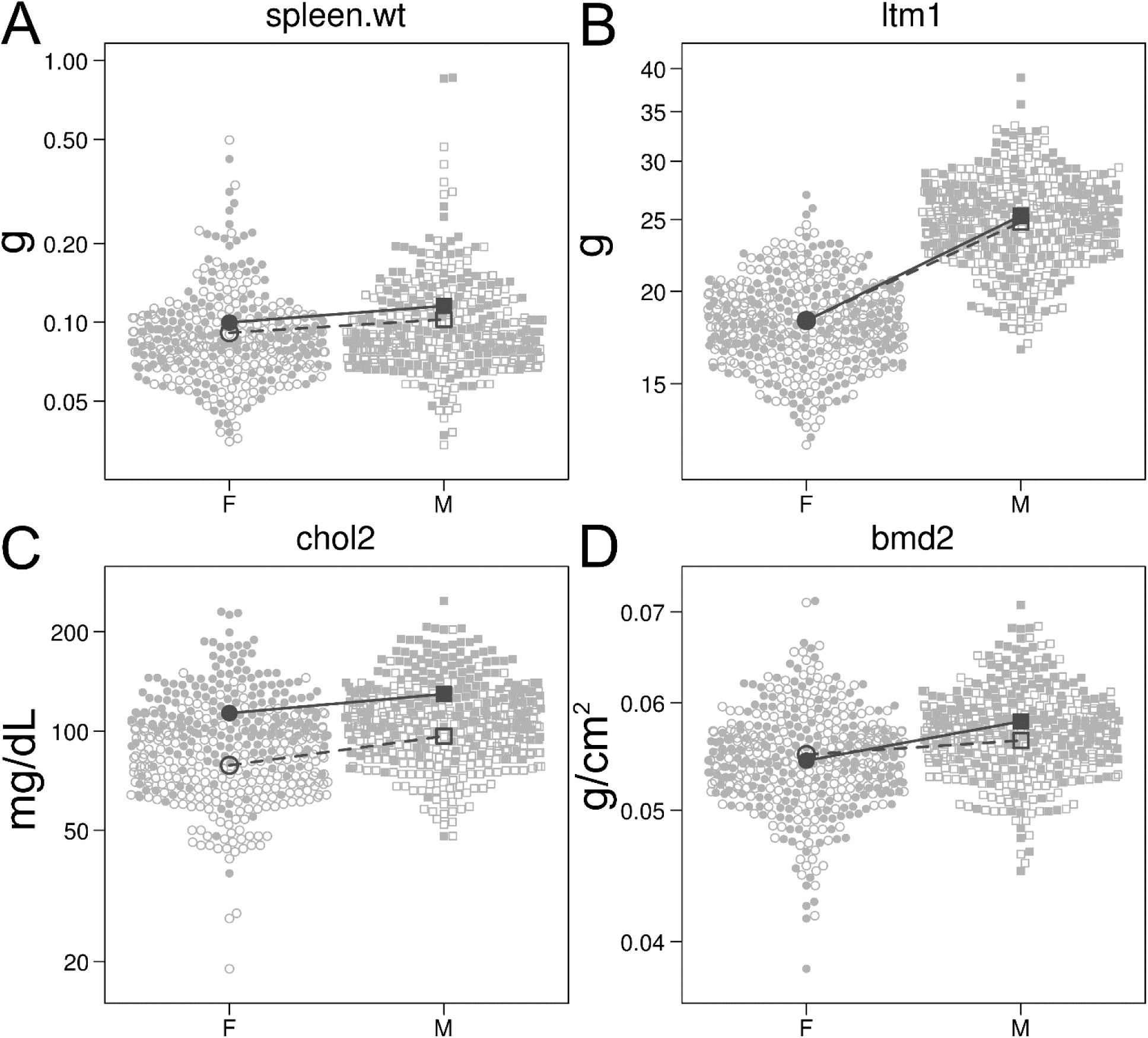
Physiological traits in the DO. Each plots shows female (circles) and male (squares) mice on chow (open symbols) and HFD (solid symbols). The open symbols connected by dashed lines show the chow diet group means. The closed symbols connected by solid lines show the HFD group means. (A) Spleen weight did not vary between sexes or diet groups. (B) Lean tissue mass at 12 weeks differed by sex and was not affected by diet. (C) Cholesterol at 19 weeks had both a significant difference between sexes and diets. (D) Bone mineral density at 21 weeks had one of the most significant sex by diet interactions, showing a greater response in males to HFD

Diet had an effect on 116 traits at an FDR ≤ 0.05. Mice on the HFD had higher mean traits values for adiponectin, weight and fat traits, %Fat, GTT.AUC, HGB, insulin, leptin, total bilirubin (TBIL), glutamate dehydrogenase (GLDH) and CHOL. Mice on the chow diet had higher blood urea nitrogen (BUN), kidney weight, MONO, NEUT, PLT, triglycerides (TG) and urine creatinine and glucose.

CHOL was higher in males as compared to females at both time points and on both diets (File S5). At eight weeks, males on the chow diet (94.1 mg/dL) had CHOL levels 20% higher than females (78.4 mg/dL). At 19 weeks, CHOL levels on the chow diet were similar to levels at eight weeks in males (94.2 mg/dL) and females (75.8 mg/dL). The HFD increased CHOL levels at both time points. At eight weeks, males on the HFD (126 mg/dL) had CHOL levels that were 20% higher than females (105 mg/dL). At 19 weeks, males (128 mg/dL) had CHOL levels that were 16.3% higher than females (110 mg/dL) (Figure 2C). At 19 weeks, the HFD increased CHOL by 45.1% compared to the chow diet in females and by 35.9% in males. CHOL levels did not change greatly between eight and 19 weeks. Female CHOL levels on the HFD increased by 4.7% from 105 mg/dL to 110 mg/dL and males increased by 1.6% from 126 mg/dL to 128 mg/dL. Therefore, the increase in CHOL levels compared to chow values was established by eight weeks in the HFD group and increased only minimally by the second time point.

There were 14 traits for which sex and diet showed a significant interaction (FDR ≤ 0.05, Figure 2D). BMD2 had one of the strongest sex by diet interactions, with mice on the chow diet having similar BMD between the sexes, but males having higher BMD than females on the HFD. This may be due to males gaining more weight and needing stronger bones to carry the weight. This is consistent with higher BW in males and the correlation of BW to BMD, such that male weight gain may require bone fortification to support increased body mass.

### Impact of Sex and Diet on Liver Transcription

We tested each of the 12,067 transcripts for the effects of sex, diet and a sex by diet interaction (File S6). We found 7,723 genes with sex effects (FDR ≤ 0.01) and 5,299 genes with significant diet effects (FDR ≤ 0.01). We found 757 genes with a significant sex by diet interaction. In order to interpret the functional relevance of these large gene lists, we searched for Gene Ontology (GO) categories that were enriched in for each effect. We identified 212 GO Biological Process (GO.BP) categories out of 2,570 in which genes were differentially expressed between the sexes (p ≤ 0.05, File S7). Of those traits affected by sex, organ regeneration (GO:00031100) was the most significant category and was higher in males. This was followed by lipid metabolism (GO:0006629) and transport (GO:0006869), which was higher in males. However, cholesterol metabolism (GO:0008203, GO:0006695) was lower in males. Fatty acid metabolism (GO:0000038 & GO:0070542) and beta-oxidation were higher in male mice along with catabolism of triglycerides (GO:0019433). Of note, male mice had higher expression of genes involved in unfolded protein responses (GO:1900103, GO:0072321), extracellular matrix disassembly (GO:0022617), and fibroblast proliferation (GO:0048146). This suggests that males experienced greater stress from unfolded protein responses, cellular remodeling and proliferation.

We identified 212 GO.BP categories that contained genes that were differentially expressed by diet (p ≤ 0.05, Supplemental File S11). For these, heat generation (GO:0031649) and energy reserve metabolism (GO:0006112) were upregulated in mice on HFD. Lipid metabolism (GO:0045834) was upregulated while lipid biosynthesis (GO:0051055), including fatty acids (GO:0006633, GO:0042761) and phospholipids (GO:0008654, GO:0015914) was down-regulated in mice fed the HFD. Lipid storage (GO:0010888), cholesterol transport (GO:0030301) and gluconeogenesis (GO:0045721) were all decreased in mice fed the HFD. Insulin secretion in response to glucose stimulation was suppressed (GO:0061179) and glucose metabolism was increased (GO:0010907). Overall, the HFD produced increases in lipid and energy metabolism while decreasing lipid biosynthesis and storage, and perturbed glucose homeostasis.

### Physiological Trait Mapping

We mapped the physiological traits using each of three models: a model in which sex and diet were additive covariates (additive model); on in which sex and diet were additive covariates and sex interacted with genotype (sex-interactive model) and one in which sex and diet were additive covariates and diet interacted with genotype (diet-interactive model).

### Additive Model

We identified 82 additive QTL with genome-wide p-values ≤ 0.05 (File S9). Of these, 39 were hematology traits, 23 were body weight or body composition traits, 14 were clinical chemistry traits, 3 were electrocardiogram traits and 3 were urinalysis traits.

Circulating cholesterol at 8 and 19 weeks (CHOL1 & CHOL2) had additive QTL on chromosome 1 at 171.37 Mb with a LOD of 13.92 for CHOL1 and 13.57 for CHOL2 (Figure 3A). The pattern of founder allele effects at this locus was similar at both time points (Figure 3B). The 129S1/SvImJ (129S1) and WSB/EiJ (WSB) alleles on the distal end of chromosome 1 were associated with higher cholesterol levels. We performed association mapping around the peak at 171.37 Mb and found that the most significant SNPs (rs587286870 & rs580179709) had founder allele patterns for which the 129S1 and WSB strains carried the minor allele (Figure 3C). When we fit the association mapping model again by including these SNPs as covariates, the maximum LOD score in the region between 170 and 175 Mb decreased to 1.72, which is well below the significance threshold. Both rs587286870 and rs580179709 are in an intron of prefoldin 2 (*Pfdn2*), which is part of a molecular chaperone complex that stabilizes unfolded proteins. It is unclear how *Pfdn2* might impact circulating cholesterol levels. However, we note that both of these SNPs are near a gene that is known to influence cholesterol levels, apolipoprotein A-II (*Apoa2*) located at 171.2 Mb. The 129S1 strain carries a private alanine to valine substitution (rs8258226) that increases cholesterol levels (Wang *et al.* 2004). The WSB strain carries a non-synonymous SNP (rs229811374) that changes a serine to an asparagine and is located six nucleotides upstream of rs8258226. If both of these SNPs increase cholesterol levels, this may explain the increase in the 129S1 and WSB alleles effects at the Apoa2 locus (Figure 3D) and the strong association with all SNPs where the minor allele occurs in both 129S1 and WSB. The peak in this region is broad and may include more than one locus and allele. When we regressed out the effects of the 129S1 and WSB alleles at the *Apoa2* locus, the maximum LOD of 5.35 occurred at 138.178 Mb on chromosome 1. We searched for other genes expressed in the liver that might influence cholesterol levels by mediating the peak with the expression of each gene (see Methods). We found six genes that reduced the LOD score by greater than six standard deviations (i.e. had a Z-score < −6, Figure 3E); inhibitor of kappaB kinase epsilon (*Ikbke*), peptidase M20 domain containing 1 (*Pm20d1*), adenosine A1 receptor (*Adora1*), coagulation factor XIII, beta (*F13b*), complement factor H-related 1 (*Cfhr1*) and cathepsin E (*Ctse*). Of these, *Cfhr1* had a Z-score of −18.6, which was lower than any other gene. The next lowest Z-score was −13 for *Ctse.* We tested whether *Ctse* would still reduce the LOD score by performing mediation analysis with *Cfhr1* in the model and found that *Ctse* still had a Z-score of −9.6. We found that *F13b* had a Z-score of −6.7 in this scan as well. The founder allele effects for *Cfhr1* (Figure 3F) show that mice carrying the PWK/PhJ (PWK) allele have higher *Cfhr1* levels and mice carrying the A/J allele have lower levels. For *Ctse,* the 129S1, WSB and NZO/HlLtJ (NZO) strains have higher *Ctse* expression (Figure 3G). For *F13b*, the WSB allele is associated with lower *F13b* expression (Figure 3H). *Cfhr1* is upregulated in the mouse retina in response to a different high fat diet (Zheng *et al.* 2015). *Ctse* deficient mice fed a high fat diet showed hypercholesterolemia, reduced body weight gain and impaired fat development compared to controls mice (Kadowaki *et al.* 2014). *F13b* is part of the coagulation cascade and has not been previously associated with cholesterol metabolism.

**Figure 3.**
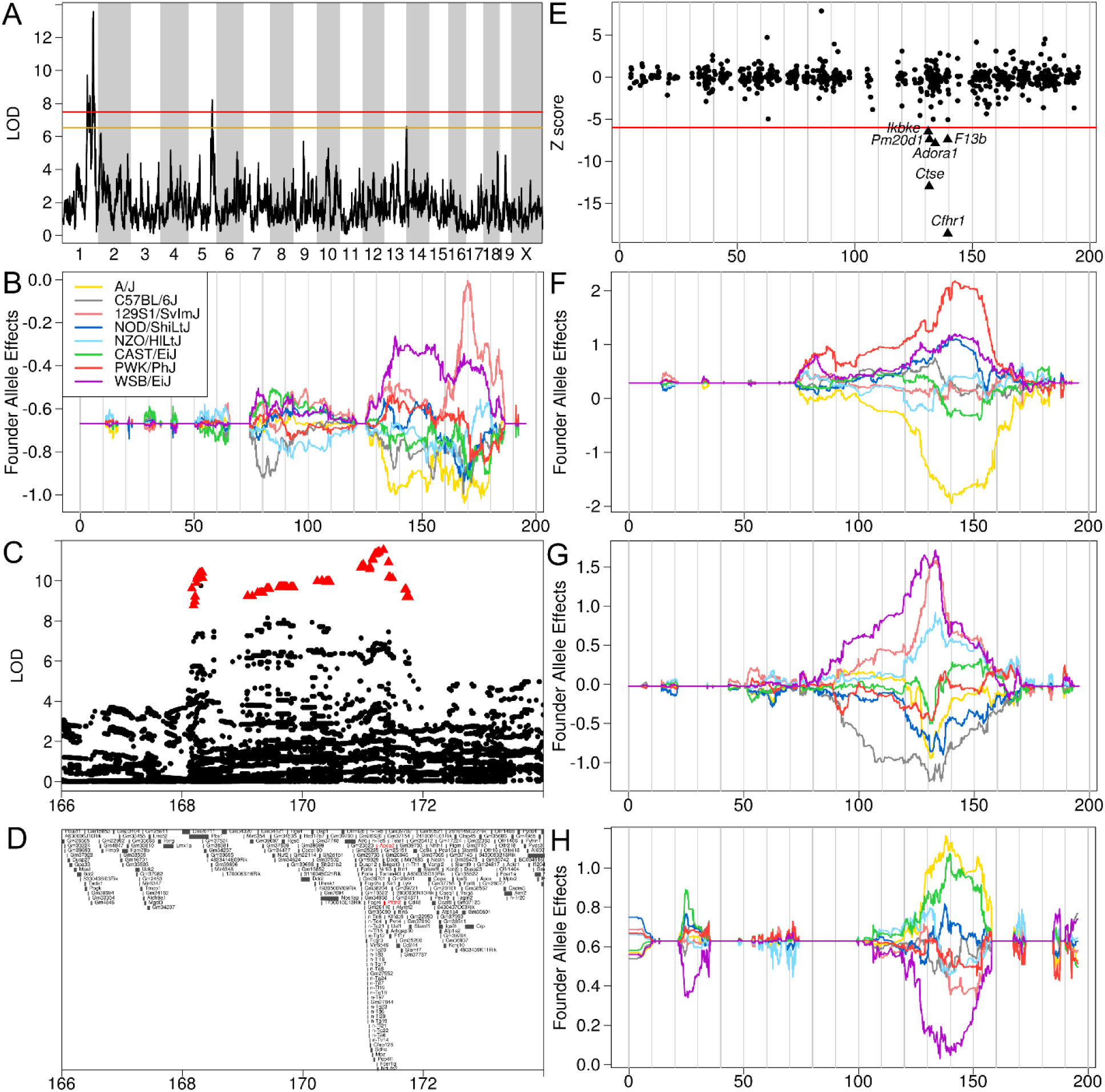
Analysis of cholesterol QTL on chromosome 1. (A) Genome scan of cholesterol at 19 weeks shows peaks on chromosomes 1 and 5. The horizontal axis shows the mouse genome and the vertical axis shows the LOD score. Red and yellow lines are the p = 0.05 and 0.2 significance levels, respectively. The horizontal axis in panels B through H shows the location in Mb on chromosome 1. (B) DO founder allele effects on chromosome 1 for cholesterol at 19 weeks (CHOL2). Each colored line represents the estimated effect of one of the founder alleles along the chromosome. (C) Association mapping near the peak on chromosome 1 shows that the SNPs for which 129S1 and WSB contribute the minor allele have the highest LOD scores (red triangles). (D) Genes in the region one chromosome 1 shown in panel C. *Apoa2* and *Pfdn2* are colored in red and are mentioned in the text. (E) Mediation analysis shows that six genes reduce the LOD score by more than 6 standard deviations. The vertical axis shows the Z-score (scaled LOD across all genes). Each point represent the Z-score (standardized LOD score reduction) for one gene in the mediation analysis. The genes are located along the horizontal axis. The red line shows Z = −6. DO founder allele effects for liver expression of (F) *Cfhr1*, (G) *Ctse* and (H) *F13b* on chromosome 1.

We found a peak for CHOL2 on chromosome 5 at 123.760 Mb for which the NZO allele was associated with higher cholesterol levels (Figure 4A). When we mediated the QTL peak at 123.76 Mb with the liver expression of each gene on chromosome 5, we found that TRAF-type zinc finger domain containing 1 (*Trafd1*) and scavenger receptor class B1 (*Scarb1*) reduce the LOD score by more than six standard deviations (Figure 4B). *Trafd1* is associated with the regulation of innate immune responses and is not known to have a function in cholesterol metabolism or transport. However, the pattern of allele effects (Figure 4C) and the mediation score suggest that it may play an unknown role.

**Figure 4.**
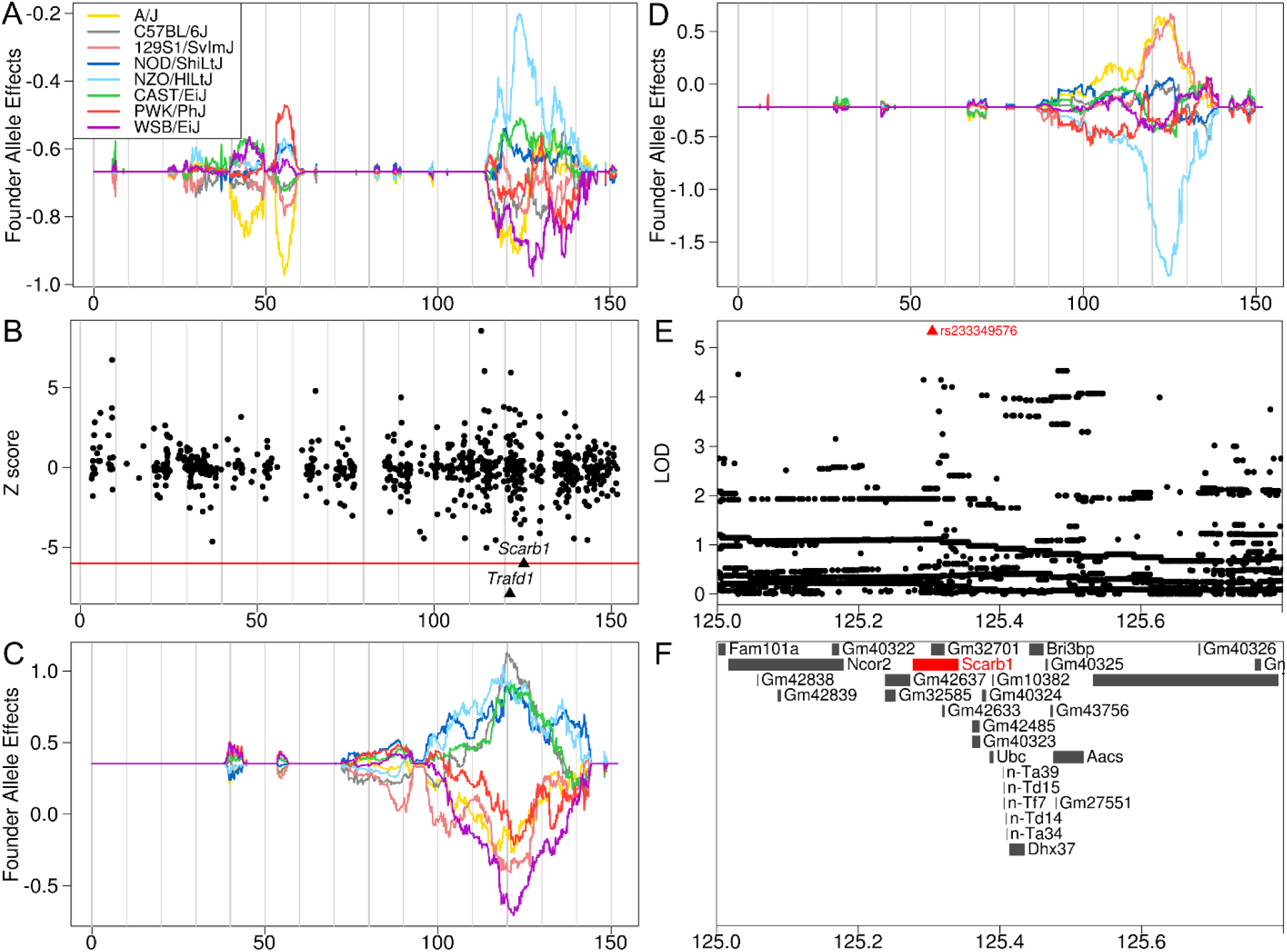
Cholesterol QTL at 19 weeks (CHOL2) on chromosome 5 suggest *Trafd1* and *Scarb1* as a candidate gene for circulating cholesterol levels. (A) Founder allele effects for CHOL2 show that the NZO allele at 123.7 Mb is associated with higher cholesterol levels. Each colored line represents the estimated effects of one founder allele. (B) Mediation analysis of the CHOL QTL on Chr 5 using liver transcripts as mediators. Each dot represents the Z-score of the LOD decrease after including one gene in the mapping model. The red line is the Z = −6 threshold. (C) Founder allele effects for liver *Trafd1* transcript levels have a similar pattern of allele effects as CHOL2. (D) Founder allele effects for liver *Scarb1* transcript levels show that DO mice with the NZO allele on chromosome 5 at 123.7 Mb have lower levels of *Scarb1*. (E & F) Association mapping in the interval near the QTL identifies a single SNP (rs233349576) that introduces a stop codon into a *Scarb1* transcript.

We repeated the mediation scan using *Trafd1* expression as a covariate and found that *Scarb1* was the only gene that reduced the LOD score by more than 6 standard deviations. DO mice carrying the NZO allele at the QTL had lower transcript levels of *Scarb1* (Figure 4D), which is consistent with the founder allele effects for CHOL2. *Scarb1* is the primary receptor for HDL-cholesterol uptake by the liver and steroidogenic tissues and is vital for reverse cholesterol transport. Targeted mutations in *Scarb1* lead to abnormal lipoprotein metabolism and increased cholesterol levels (Mohr *et al.* 2004). Liver-specific reduction in Scarb1 expression as a result of an ENU-induced point mutation has also been reported, in which mice exhibit 70% higher plasma HDL-cholesterol levels due to reduced HDL selective uptake (Stylianou *et al.* 2009). *Scarb1* was proposed as a candidate gene for hypercholesterolemia in an intercross between NZB/B1NJ and SM/J (Pitman *et al.* 2002), but the authors found no sequence, mRNA or protein differences. However, whole-genome sequencing of NZO has revealed a stop gain mutation (rs233349576, Figure 4E & F) at residue 37 in ENSMUST00000137783. This may produce an incomplete protein and may alter its function.

### QTL that Interact with Diet

There were 12 QTL with p-values ≤ 0.05 for which genotype interacted with diet (File S9), including 6 clinical chemistry traits, heart rate, lymphocyte counts, urinary creatinine, body weight and body length. Cholesterol at 8 weeks (CHOL1) had a QTL that interacted with diet on chromosome 10 at 21.99 Mb with a LOD of 10.6 (p ≤ 0.001, Figure 5A). Mice carrying the NOD allele on the HFD had higher cholesterol levels. We mediated the QTL peak with all liver transcripts and found that *E030030I06Rik* decreased the LOD by more than six standard deviations. *E030030I06Rik* has a local eQTL on chromosome 10 at the same location. Association mapping near the QTL peak produced significant associations with one gene, *Gm20125,* which is a gene model with no known function.

**Figure 5.**
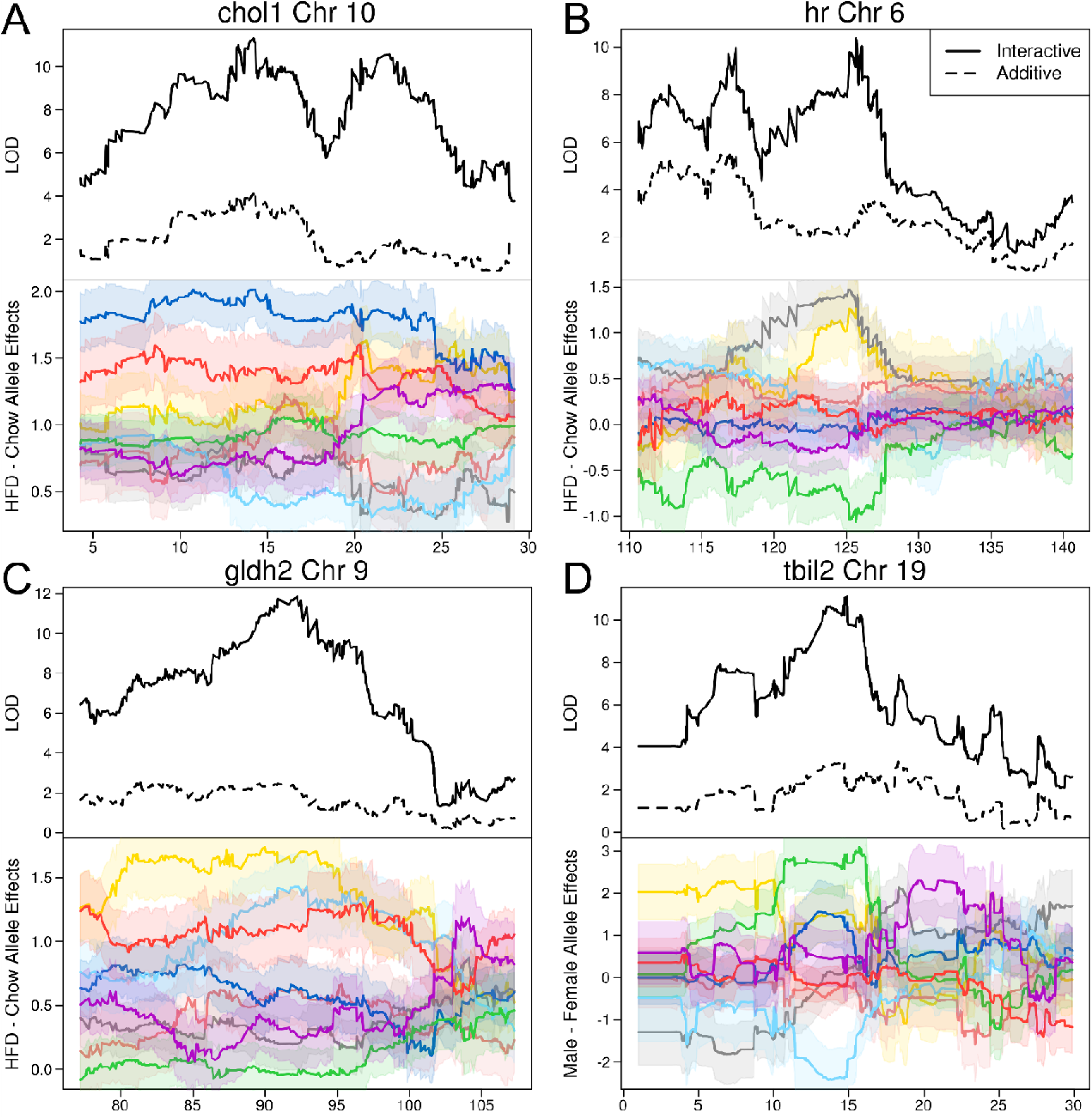
Sex- and diet-interactive QTL plots. Each plot has two panels. The top panel shows the additive LOD (dashed line) and the interactive LOD (solid line). The lower panel shows the difference between the interactive and additive founder allele effects, with standard errors in light shading. Each founder is represented by a separate color: A/J yellow, C57BL/6J grey, 129S1/SvImJ pink, NOD/ShiLtJ cyan, NZO/HlLtJ blue, CAST/EiJ green, PWK/PhJ red and WSB/EiJ purple. (A) Cholesterol at eight weeks had a diet-interactive QTL on chromosome 10. (B) Heart rate had a diet-interactive QTL on chromosome 6. (C) Glutamate dehydrogenase at 19 weeks had a diet-interactive QTL on chromosome 9. (D) Bilirubin at 19 weeks had a sex-interactive QTL on chromosome 10.

Heart rate at 13 weeks (HR) had a QTL that interacted with diet on chromosome 6 at 125.63 MB with a LOD of 10.4 (p ≤ 0.001, Figure 5B). Mice carrying the A/J or C57BL/6J allele on the HFD had higher HR than mice carrying the CAST allele. We did not perform mediation analysis because we do not have heart transcript information on these mice. Association mapping near the peak produced two SNPs (rs48596855, rs38346309) in the introns of anoctamin 2 (*Ano2*), a calcium activated chloride channel, a class of genes that may play a role in cardiac function (Guo *et al.* 2008; Hartzell *et al.* 2009). Another gene, potassium voltage-gated channel, shaker-related 1, (*Kcna1*), is located 1 Mb distal from the peak SNPs and has been associated with changes in heart rate (Glasscock *et al.* 2010).

Glutamate dehydrogenase, a marker of liver injury, at 19 weeks (GLDH2) had a QTL that interacted with diet on chromosome 9 at 92.19 Mb with a LOD of 11.86 (p ≤ 0.001, Figure 5C). Mice carrying the A/J, NZO or PWK alleles on the HFD had higher GLDH levels. There were no genes that had mediation Z-scores less than −6 near the QTL peak. When we performed association mapping near the peak, we found four transcripts with intronic SNPs and LOD scores over 7; *Gm29478, 1700057G04Rik*, phospholipid scramblase 4 (*Plscr4*), and procollagen lysine, 2-oxoglutarate 5-dioxygenase 2 (*Plod2*). Both *Plscr4* and Plod2 had local eQTL on chromosome 9. *Plscr4* is a membrane protein that is involved in the organization of phospholipids and interacts with the CD4 receptor of T lymphocytes (Py *et al.* 2009). It is up-regulated with HFD feeding (Song *et al.* 2012). *Plod2* hydroxylates lysine residues and is involved in remodeling of the extracellular matrix (Gilkes *et al.* 2013) and fibrosis (van der Slot *et al.* 2003). Under hypoxic conditions, *Plod2* is expressed in hepatocellular carcinoma and is correlated with tumor size and macroscopic intrahepatic metastasis. It is a prognostic factor for disease-free survival (Noda *et al.* 2012).

### QTL that Interact with Sex

There were 9 QTL with p-values ≤ 0.05 for which genotype interacted with sex (File S9), including 5 clinical chemistry traits and 4 hematology traits. Blood urea nitrogen at 19 weeks (BUN2) had a QTL that interacted with sex on chromosome 10 at 95.85 Mb with a LOD of 11.7 (p ≤ 0.001). Males carrying the 129S1, C57BL/6J and WSB alleles were associated with higher BUN and females carrying the PWK allele with lower BUN. We did not perform mediation analysis because we do not have kidney transcript information on these mice. Total bilirubin at 19 weeks (TBIL2) had a QTL that interacted with sex on chromosome 19 at 14.89 Mb with a LOD of 11.1 (p ≤ 0.001, Figure 5D). Females carrying the NZO allele had higher bilirubin and males carrying the CAST allele had lower bilirubin. Mediation analysis using liver gene expression did not reveal a candidate gene. The most significant SNPs in the association mapping on chromosome 19 were near five transcripts: Gm8630, Gm31441, Gm37997, Gm26026 and transducin-like enhancer of split 4 (*Tle4*). None of the gene models had a QTL. *Tle4* is a transcriptional corepressor factor that regulates mouse hematopoiesis and bone development (Wheat *et al.* 2014), and has also been used for histological application as a podocyte nuclear marker in glomeruli (Venkatareddy *et al.* 2014).

### Liver Expression QTL Mapping

We performed linkage mapping on 12,067 liver genes and identified additive QTL (File S10), QTL that interact with sex (File S11) and QTL that interact with diet (File S12). We mapped local and distant eQTL using an additive linkage model, and two models in which sex or diet interacts with genotype. We plotted the location of significant QTL versus gene location for each model and found that the additive and sex-interactive models produced local eQTL (Figure 6A & B) and the diet-interactive model did not (Figure 6C). At a significance threshold of 0.05, we found that 8,127 local eQTL out of 9,754 total (83.3%) in the additive model, 332 out of 532 (62.4%) in the sex-interactive model and 23 out of 219 (10.5%) in the diet-interactive model. We have provided an on-line visualization tool at http://churchill-lab.jax.org/qtl/svenson/DO478/.

**Figure 6.**
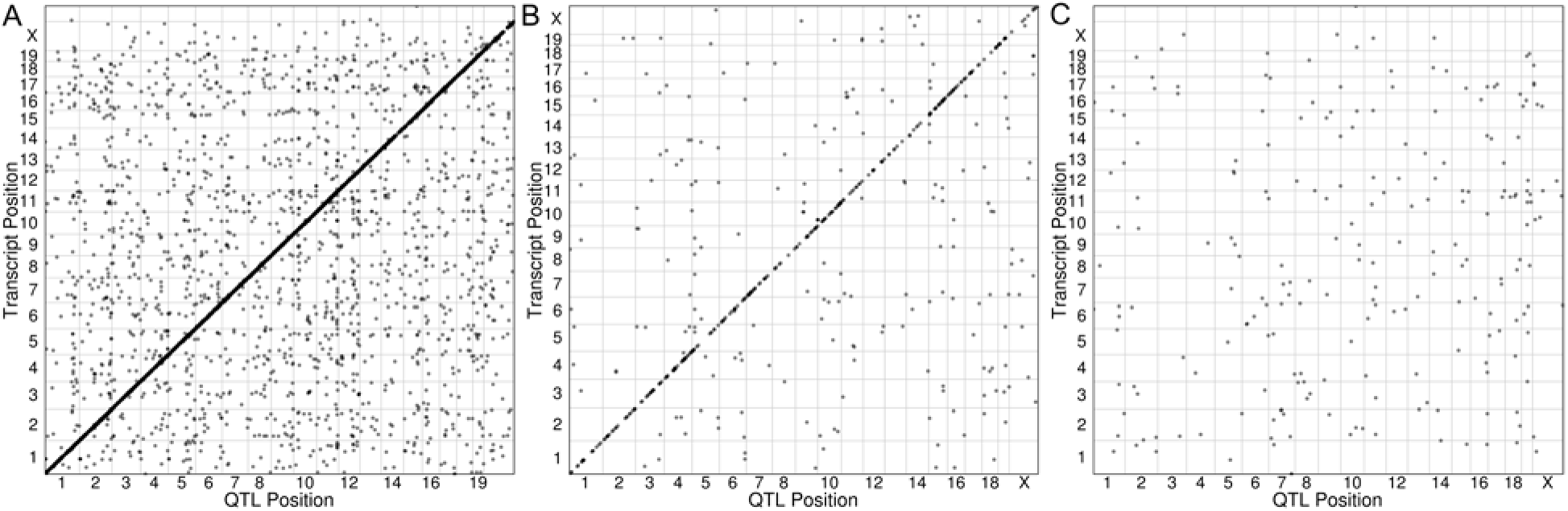
Liver expression QTL maps generated from an (A) additive model (File S10), (B) a model in which sex interacts with genotype (File S11), or (C) a model in which diet interacts with genotype (File S12). Each dot represents the location of a QTL peak for one gene above the p = 0.05 threshold. Each panel plots the QTL position on the horizontal axis and the transcript position on the vertical axis.

### XO Females

XO females have been previously reported in high numbers in DO mice (Chesler *et al.* 2016). In order to search for XO females, we plotted the liver expression of *Xist* as an indicator of X chromosome gene expression versus *Ddx3y* as a marker of Y chromosome gene expression (Figure 7). As expected, females that were dizygous for the X chromosome had high *Xist* expression and low *Ddx3y* expression while males had high *Ddx3y* expression and lower *Xist* expression. There were two females (out of 244, 0.82%) that had low *Xist* expression, consistent with hemizygosity on the X chromosome, and low *Ddx3y* expression and these samples are XO females. In humans, Turner Syndrome describes females with the XO genotype and is characterized by short stature, a propensity for ovarian dysfunction and infertility, and heart defects. The two XO females in our study, one fed chow and the other fed the HFD, had very different phenotypes and were not outliers for any particular trait. One of them, however, was especially resistant to weight gain on HFD, gaining only 5 grams of body weight over the course of the study, and died 4 weeks before the scheduled end of the study.

**Figure 7.**
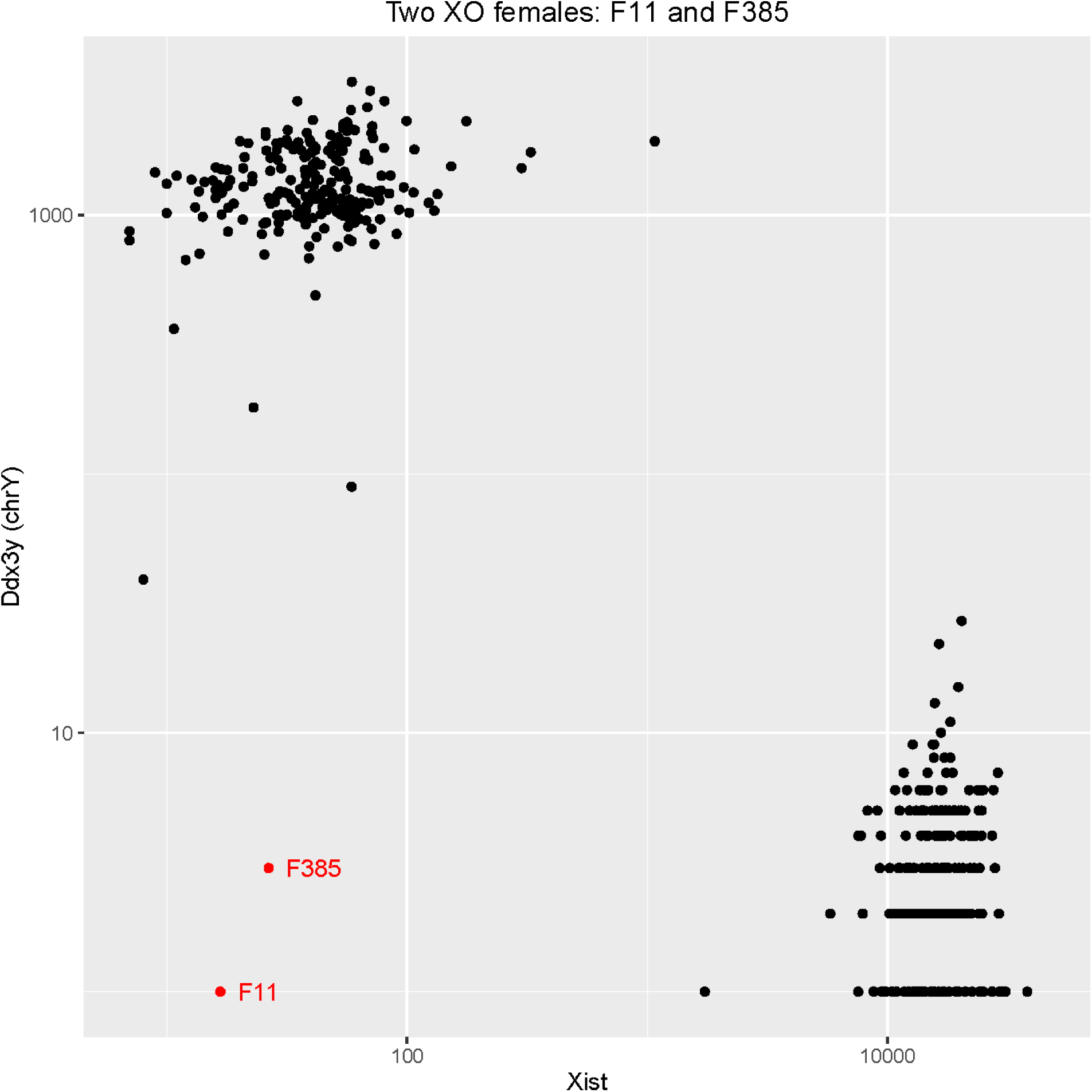
XO females in the DO population. For each mouse, we plotted the untransformed expression of *Xist* versus *Ddx3y* and found two XO females (in red). Females have high *Xist* expression and low *Ddx3y* expression. Males have low *Xist* expression and high *Ddx3y* expression.

## DISCUSSION

Multi-parent populations are excellent tools for studying the effects of genetic diversity on phenotypic variation because they offer increased genetic diversity, high minor allele frequencies and fine recombination block structure. The large number of variants leads to perturbation of genes throughout the genome and the high minor allele frequency produces high power to detect the effects of these polymorphisms. The fine recombination block structure leads to fine mapping resolution to identify loci that contain a manageable number of candidate genes. These loci may contain genetic variants that alter either protein structure or transcript levels or both. In fact, many loci in multi-parent crosses may be caused by more than one genetic variant and disentangling the signal from these different alleles is a complex process. For the cholesterol loci on chromosomes 1 and 5, we combined association mapping using imputed SNPs with mediation analysis using liver transcripts and we identified candidate genes using both missense SNPs and transcript levels. This approach is broadly applicable in the DO and other multi-parent populations, but requires transcriptional profiling in a relevant tissue.

CHOL levels were associated with two loci on chromosomes 1 and 5. Variation at these loci affected mice of both sexes and on both diets. In contrast, CHOL at eight weeks had a QTL on chromosome 10 with effects that were modified by diet. The HFD increased CHOL levels by 35.9% in males and 45.1% in females, indicating that diet increases CHOL levels in most mice. However, DO mice carrying the NOD allele on chromosome 10 have higher CHOL levels than mice carrying other alleles at the same locus. The locus covered several Mb and may contain more than one polymorphism that affects CHOL levels, independent of diet.

Association mapping on chromosome 1 produced a set of SNPs with high LOD scores for which 129S1 and WSB contributed the minor allele. Initially, we expected to find SNPs with this allele pattern that alter the protein structure or expression levels of some gene. However, in this case, we believe that two closely located SNPs in *Apoa2*, each of which is private to either 129S1 or WSB, are influencing CHOL levels. The SNP in 129S1 (rs8258226) has been shown to increase CHOL levels and we hypothesize that the SNP in WSB (rs229811374), which is 2 residues away from rs8258226, may also increase CHOL levels. This highlights the complexity of analyses in multi-parent crosses. Had we not known of any candidate genes in this region, we may have been led to consider other genes that are unrelated to CHOL metabolism.

Mediation analysis identified several candidate genes on chromosome 1. By its nature, mediation analysis is a hypothesis generating analysis. All of the genes, *Cfhr1*, *Ctse* and *F13b* are within 10 Mb of each other. Of these, *Ctse* is the only gene that had been previously associated with hypercholesterolemia (Zheng *et al.* 2015). The situation is similar for the CHOL peak on chromosome 5. *Scarb1* has been associated with differences in CHOL levels, but *Trafd1* is a new candidate gene that may have effects of CHOL independent of *Scarb1*. These findings suggest that some strong associations that appear in MAGIC populations may be due to multiple, tightly linked polymorphisms. If this is true, more sophisticated, multi-locus approaches to candidate gene selection will be needed to find causal genes. In this study, we suggest that both association mapping and mediation analysis with transcript levels improve the reliability of candidate gene selection because it allows investigators to search for polymorphisms that affect protein structure or transcript levels.

When we performed liver eQTL mapping, we found local eQTL for both the additive model and the sex-interactive model. This suggests that local polymorphisms in regulatory elements modulate transcript levels constitutively in both sexes and differentially by sex. However, when we mapped liver transcript levels using a model in which diet interacts with genotype, we found very few local eQTL. This suggests that the response to diet is less influenced by the interaction of diet with local polymorphisms affecting transcript levels, we speculate that distant loci are acting through non-transcriptional mechanisms such as interactions between proteins (Chick *et al.* 2016).

Our analysis of this large study using DO mice includes a novel multi-tiered approach that has identified plausible candidate genes underlying physiological traits, has extended to considering effects of transcription, and provides compelling evidence for further investigation of the role of novel candidates in driving metabolic traits. We present an analysis of the complex interplay of sex and diet and how these factors influence important inter-individual variation in outcome. We found that diet increases many traits related to body size and composition. Interestingly, we observed that a high fat, high sucrose diet increased the variance of liver gene expression, suggesting that genotype influences the range of responses to diet. We mapped loci for multiple traits but focused on CHOL to demonstrate the complexity of the underlying genetic loci. We note that both association mapping and mediation analysis using liver transcript data help to dissect the causal alleles underlying mapping peaks. Finally, we observe that changes in liver transcription in response to diet are not primarily altered by local genetic variants. This suggests that other molecular measurements, such as protein or metabolite levels may be more useful in determining the effects of diet on organisms. We have made the phenotype and genotype data fully available to the public and have released an interactive viewer to allow the reader to explore the liver expression QTL data.

## ACKNOWLEDGEMENTS

This work was funded by P50GM076468 and R01GM070683 to G.A.C. from the National Institute of General Medical Sciences.

